# Thalamocortical And Intracortical Inputs Differentiate Layer-Specific Mouse Auditory Corticocollicular Neurons

**DOI:** 10.1101/288431

**Authors:** Bernard J. Slater, Stacy K. Sons, Daniel A. Llano

**Affiliations:** Neuroscience Program, University of Illinois Urbana-Champaign, Urbana, IL; Department of Molecular and Integrative Physiology, University of Illinois Urbana-Champaign, Urbana, IL; Beckman Institute for Advanced Science and Technology, Urbana, IL

## Abstract

Long-range descending projections from the auditory cortex play key roles in shaping response properties in the inferior colliculus. The auditory corticocollicular projection is massive and heterogeneous, with axons emanating from cortical layers 5 and 6, and plays a key role in directing plastic changes in the inferior colliculus. However, little is known about the cortical and thalamic networks within which corticocollicular neurons are embedded. Here, laser scanning photostimulation glutamate uncaging and photoactivation of channelrhodopsin-2 were used to probe the local and long-range network differences between pre-identified mouse layer 5 and layer 6 auditory corticocollicular neurons in vitro. Layer 5 corticocollicular neurons were found to vertically integrate supragranular excitatory and inhibitory input to a substantially greater degree than their layer 6 counterparts. In addition, all layer 5 corticocollicular neurons received direct and large thalamic inputs from channelrhodopsin-2 labeled thalamocortical fibers whereas such inputs were less common in layer 6 corticocollicular neurons. Finally, a new low calcium/synaptic blockade approach to separate direct from indirect inputs using laser photostimulation was validated. These data demonstrate that layer 5 and 6 corticocollicular neurons receive distinct sets of cortical and thalamic inputs, supporting the hypothesis that they have divergent roles in modulating the inferior colliculus. Furthermore, the direct connection between the auditory thalamus and layer 5 corticocollicular neurons reveals a novel and rapid link connecting ascending and descending pathways.

## Introduction

As we navigate the environment, we are presented with an array of complex sounds which must be selected, enhanced or suppressed to guide behavior. Descending projections from the cortex are responsible for much of the shaping of ascending auditory information, particularly at the level of the inferior colliculus, which is the main integration site for acoustic information prior to reaching the forebrain (for reviews see (Winer, 2006; Suga, 2008; Bajo and King, 2012; Stebbings et al., 2014)). However, despite the number of studies examining the impact of this projection on midbrain response properties, little is known about corticocollicular circuitry. At the level of the cortex, neurons from two layers project to the inferior colliculus: those from layer 5 and a smaller proportion (~10-20% in mouse) that are found in deep layer 6 (Games and Winer, 1988; Künzle, 1995; Coomes et al., 2005; Bajo et al., 2007; Schofield and Motts, 2009; Slater et al., 2013). Layer 5 corticocollicular neurons are large pyramidal cells with a thick apical dendrite projecting toward layer 1, and many of these cells rhythmically burst when depolarized. Layer 6 corticocollicular cells, in contrast, are non-pyramidal, regular-spiking cells with elongated somata and thin, but densely branching dendrites (Slater et al., 2013; Zurita et al., 2017). Therefore, similar to the corticothalamic system where layer 5 and layer 6 neurons have distinct roles in modulating the thalamus (Ojima, 1994; Reichova and Sherman, 2004; Llano and Sherman, 2009; Theyel et al., 2010), there are two major types of cells comprising the corticocollicular projection which may be responsible for the diverse collicular responses observed after manipulations of this pathway.

To better understand the mechanisms by which the auditory cortex modulates the inferior colliculus, it is important to understand how corticocollicular neurons are integrated into local cortical and thalamocortical networks. Previous studies of layer 5 cortical neurons have shown that these cells receive both local excitatory and inhibitory cortical inputs from near the soma as well as from upper layers (Schubert et al., 2001; Thomson et al., 2002; Schubert et al., 2006; Llano and Sherman, 2009; Hooks et al., 2013; Zarrinpar and Callaway, 2016). This distribution of inputs is in stark contrast to the inputs to layer 6 corticothalamic cells which receive the majority of their excitatory and inhibitory input from layer 6 (Llano and Sherman, 2009). With respect to thalamic input, while the canonical view is that the thalamus provides input to layer 4, thalamocortical fibers also have been shown to contact other layers, including layer 5 and layer 6 neurons (Huang and Winer, 2000; Zhao et al., 2009; Yang et al., 2014; Ji et al., 2015). In addition, direct thalamic input to the infragranular cortical layers has been shown to result in suprathreshold responses, particularly in layer 5 intrinsically bursting neurons (Constantinople and Bruno, 2013). These layer 5 neurons send their axonal projections to subcortical nuclei, providing a rapid route to connect the ascending and descending systems. However, the presence of a direct thalamic projection to auditory corticocollicular neurons has not yet been investigated. Such a pathway could be a substrate for nearly “online” modulation of ascending signals, where some of the input is relayed to lower centers and is minimally processed in the cortex prior to transmission.

Therefore, we examined the local cortical and thalamocortical networks within which layer 5 and layer 6 corticocollicular neurons are embedded. To do this, we employed a range of optical stimulation approaches and validated an approach to separate synaptic versus direct glutamate stimulation of cortical neurons. The results of this work demonstrate a previously unknown degree of synaptic heterogeneity influencing the auditory corticocollicular system, and may account for the myriad effects of cortical stimulation on the auditory midbrain (Mitani et al., 1983; Zhang et al., 1997; Gao and Suga, 2000; Ma and Suga, 2001; Yan and Ehret, 2001; Nakamoto et al., 2008; Bajo et al., 2010; Nakamoto et al., 2010; Robinson et al., 2016).

## Methods

### Animal Preparation

All procedures were approved by the Institutional Animal Care and Use Committee at the University of Illinois. All animals were housed in animal care facilities approved by the American Association for Assessment and Accreditation of Laboratory Animal Care. BALB/c (30-60 days of age) mice were bred in house. For injections of tracers, mice were anesthetized with an intraperitoneal injection of ketamine hydrochloride (100 mg/kg) and xylazine (3 mg/kg) and then carefully placed in a stereotaxic apparatus to avoid damage to peripheral auditory structures. Lidocaine (1%) was injected subcutaneously at incision sites prior to surgery as a supplement to anesthesia. Injection targets in the inferior colliculus (Figure 1A), and medial geniculate body (MGB, Figure 1D) were localized using stereotactic coordinates (for inferior colliculus injections, 0.75 mm caudal from lambda, 0.75 mm lateral to midline, and 0.5-1 mm depth from the dorsal surface; for MGB injections, 3.18 mm caudal from bregma, 2.0 mm lateral to the midline, and 3.2 mm from the dorsal surface). No attempt was made to isolate injections to individual subdivisions of the target structures. For all animals, micropipettes (tip diameter 10 μm) were filled with 10-25 nL of latex microspheres (Lumafluor Retrobeads, Durham, NC) and injected into the inferior colliculus over 5-10 minutes using a Nanoliter 2000 injection system (World Precision Instruments, Sarasota, FL). To allow for retrograde transport of the beads from the inferior colliculus (Figure 1B) to layer 5 and layer 6 corticocollicular neurons in the auditory cortex (Figure 1C), animals were euthanized >3 days after the injection. For animals receiving MGB injections, at least 2 weeks prior to inferior colliculus injections, animals were injected with 10-25 nL of the AAV2 construct of AAV-CaMKIIa-hChR2(H134R)-mCherry, UNC Vector Core, Chapel Hill, NC) virus at 4.55×10^12^ viral genomes/mL diluted in 1% Polybrene in phosphate-buffered saline (PBS) into the MGB (Figure 1E) resulting in robust expression in thalamocortical afferents (Figure 1F). Although the retrogradely-labeled neurons (labeled with red retrobeads) and thalamocortical afferents (labeled with mCherry) fluoresce using similar wavelengths, they are easily distinguishable under high magnification. Other combinations of fluorophores (e.g., use of green retrobeads) would have caused high levels of activation of labeled thalamocortical afferents when using blue light to visualize the labeled neurons.

**Figure 1:**
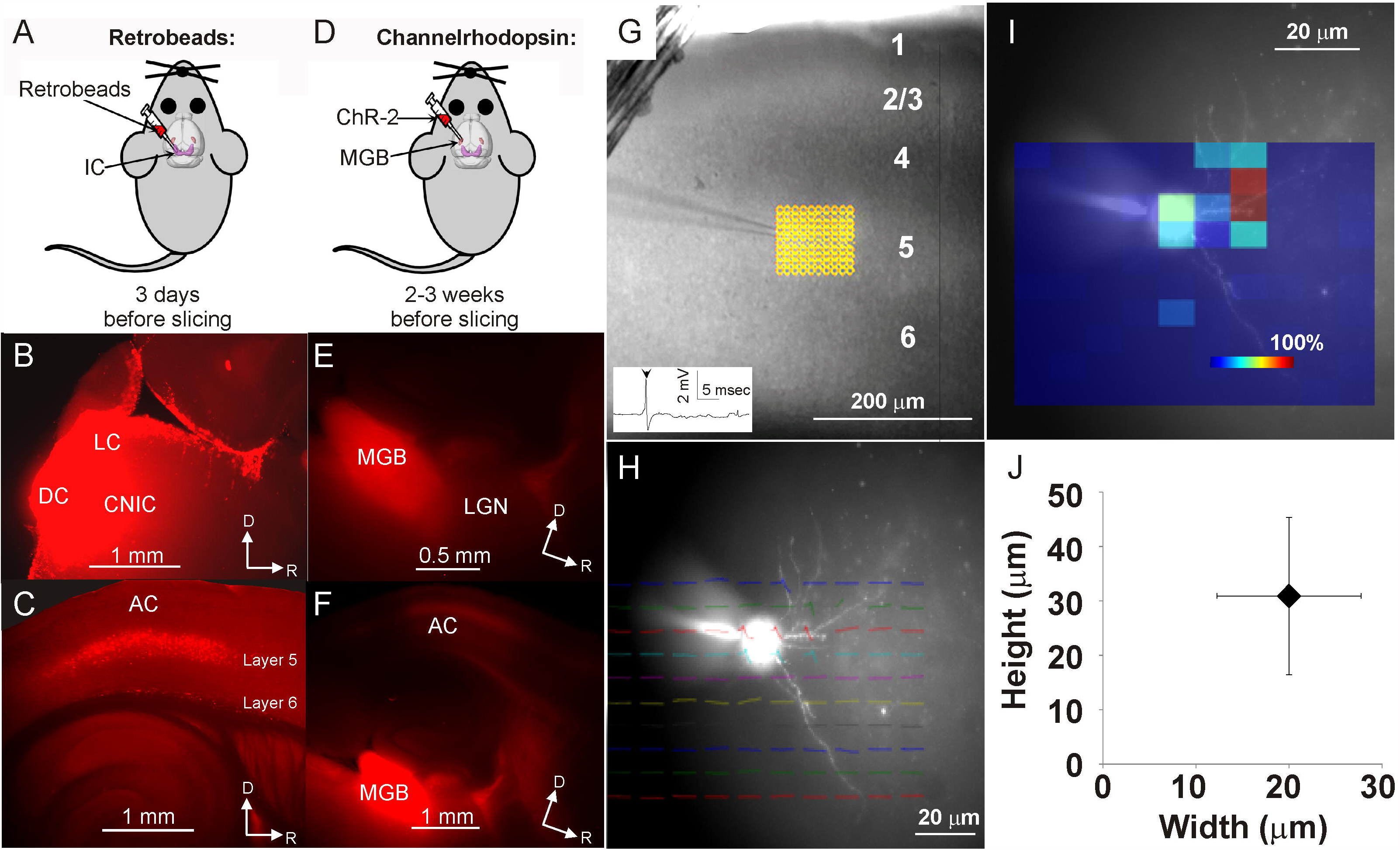
Overview of injections and cell-attached recordings. (A) Latex beads were injected into the inferior colliculus at least 3 days prior to brain slicing. (B) Inferior colliculus injection site covering portions of the lateral cortex (LC), dorsal cortex (DC) and central nucleus of the inferior colliculus (CNIC). (C) Layer 5 and layer 6 auditory corticocollicular neurons are filled with latex beads after retrograde transport. (D) Channelrhodopsin-2 AAV was injected into the MGB at 2-3 weeks prior to preparation of brain slices. (E) Injection site of AAV-CaMKIIa-hChR2(H134R)-mCherry showing the labeled MGB region and radiating labeled axons. (F) After 2-3 weeks channelrhodopsin-2 is robustly expressed in the MGB with afferents projecting to the auditory cortex. (G) Unlabeled neurons across layers 2 through 6 of the auditory cortex were recorded from using a cell-attached approach. A tight 10×10 grid of points with 10 µm spacing was centered over the cell and glutamate was uncaged in successive non-neighboring points. Inset: Spiking activity was determined visually (arrowhead). (H) After recording, the cell membrane seal is broken to permit entry of fluorescent Alexa hydrazide to view cell morphology. Shown here are the unshuffled traces, plotted over the filled dendritic processes of the cell. (I) Heatmap indicating the likelihood of eliciting a spike at each site around the neuron. (J) Average width vs. average height of the excitation profiles (n=11 cells) with standard deviation error bars. AC = auditory cortex. LGN = lateral geniculate nucleus. D=dorsal. R=rostral.

### Thalamocortical Slices

To obtain brain slices, each animal was deeply anesthetized by intraperitoneal injection of ketamine hydrochloride (100 mg/kg) and xylazine (3 mg/kg) then transcardially perfused with an ice-cold high-sucrose cutting solution (in mM: 206 sucrose, 10.0 MgCl_2_, 11.0 glucose, 1.25 NaH_2_PO_4_, 26 NaHCO_3_, 0.5 CaCl_2_, 2.5 KCl, pH 7.4) following which the brain was quickly removed. A modified version of the auditory thalamocortical slice (Cruikshank et al., 2002) was used for these experiments. As described previously (Llano et al., 2014; Slater et al., 2015), the inferior colliculus was retained in this slice to allow visualization of the colliculus injection site (Figure 1B). Slices were cut using a vibrating tissue slicer (Leica VT1000s, Buffalo Grove, IL) and transferred to a holding chamber containing oxygenated incubation artificial cerebrospinal fluid (aCSF) (in mM: 126 NaCl, 3.0 MgCl_2_, 10.0 glucose, 1.25 NaH_2_PO_4_, 26 NaHCO_3_, 1.0 CaCl_2_, 2.5 KCl, pH 7.4) at 32°C for 1 hour prior to recording. Given this orientation of slice, which is angled at about 15 degrees from the horizontal plane, the terms “dorsal” and “lateral” are somewhat ambiguous. For consistency, for the current report, “dorsal” refers to regions of the slice closer to the pial surface over the auditory cortex.

### Electrophysiological Recording

Whole-cell and cell-attached recordings were obtained using a visualized slice electrophysiology setup, outfitted with infrared-differential interference contrast optics and fluorescence, and performed at 22°C. The Multiclamp 700B amplifier (Axon Instruments, Sunnyvale, CA) and pClamp software (Molecular Devices, Sunnyvale, CA) were used for data acquisition, sampled at 20 kHz. Borosilicate glass capillary tubes were pulled to obtain recording pipettes. These pipettes were then filled with recording solution (for cell-attached recordings: in mM: 117 K-gluconate, 13 KCl, 1.0 MgCl_2_, 0.07 CaCl_2_, 0.1 ethyleneglycol-bis(2-aminoethylether)-N,N,N⍰, N⍰-tetra acetic acid, 10.0 4-(2-hydroxyethyl)-1-piperazineethanesulfonic acid, 2.0 Na-ATP, 0.4 Na-GTP, 0.01 Alexa 568 or 488 hydrazide, and 0.5% biocytin, pH 7.3 and intracellular recordings: in mM 117.0 CsOH, 117.0 Gluconic acid, 11.0 CsCl, 1.0 MgCl_2_*6H_2_0, 0.07 CaCl_2_, 11.0 EGTA, 10.0 HEPES). Laser stimulation was done with slices bathed in aCSF (in mM: 126 NaCl, 2.0 MgCl_2_, 10.0 glucose, 1.25 NaH_2_PO_4_, 26 NaHCO_3_, 2.0 CaCl_2_, 2.5 KCl, and 0.05 APV, at pH 7.4 with 150 µM MNI-glutamate (Tocris Biosciences, Ellisville, MO). APV (Tocris Biosciences, Bristol, United Kingdom), a selective competitive inhibitor of the N-Methyl D-Aspartate (NMDA) receptor, was used to limit recurrent excitation, therefore limiting excitation to monosynaptically driven currents (Shepherd, 2012).

### Photostimulation for cell-attached recordings

Recordings of laser-driven spikes were done after an observed increase in series resistance to approximately one gigaOhm. A UV laser (355nm wavelength, frequency-tripled Nd:YVO4, 100-kHz pulse repetition rate; DPSS Lasers, San Jose, CA) was used for these experiments. Laser power was adjusted using an acousto-optical modulator (Gooch and Housego, Ilminster, United Kingdom). The laser is directed into the side port of an Olympus microscope (BX51WI) using a series of UV-enhanced aluminum mirrors (Thorlabs, Newton, NJ) and a pair of mirror galvanometers (Cambridge Technology, Cambridge, MA) and is reflected off of a 400 nm long pass dichroic mirror to allow for laser stimulation as well as visualization of fluorescently labeled cells. The laser beam is focused onto the brain slice with a low-magnification objective (4× 0.13NA infinity corrected Plan, Olympus) and used to uncage glutamate at 15.8 mW, measured with a thermal sensor power meter (PM160T, Thorlabs). One msec pulses controlled by the acousto-optical modulator were used to obtain a series of traces corresponding to glutamatergic responses at each point. For cell-attached recordings used to classify the excitation profile of auditory cortical neurons, a 10×10 grid with adjacent rows and columns spaced 10 µm apart was used. Glutamate uncaging was done at successive non-neighbor points to prevent depletion of the caged glutamate, glutamate toxicity, and habituation at the probed synapses. Spikes were visually identified and assigned to points in the 10×10 maps; four runs were used to record the neuronal activations for each cortical layer. Spikes were considered to be reliably driven by laser input if they occurred within 5 msec after the laser pulse and in at least half of the repeated 10×10 maps at a given point.

### Photostimulation for corticocollicular mapping

Individual labeled corticocollicular neurons were identified using the presence of fluorescent latex beads (Lumafluor Retrobeads, Durham, NC) identified by fluorescence optics (Olympus filter set U-MWG2, excitation 510 – 550 nm, dichroic 570 nm, and emission 590 nm long pass filter) using a 200 W metal arc lamp (Prior, Rockland, MA). To obtain input maps, once a cell was successfully patched, MNI-glutamate was introduced into the circulating aCSF at 150 µM. A 30×30 grid of points with 25 µm spacing between adjacent rows and columns was then placed over the cortex surrounding the corticocollicular cell using the Prairie View software (Bruker, Billerica, MA). Each grid of laser stimulation points was run at least three times, first held at +10 mV to record inhibitory post-synaptic currents, next while the neuron was held at - 60mV to measure excitatory currents, and then the bathing solution was switched to a low calcium aCSF (in mM: 126 NaCl, 4.0 MgCl_2_, 10.0 glucose, 1.25 NaH_2_PO_4_, 26 NaHCO_3_, 0.01 CaCl_2_, 2.5 KCl, 0.05 APV, at pH 7.4), which limits synaptic transmission, while the neuron was again held at −60mV to record direct stimulation of the recorded neuron. QX-314 (Tocris Biosciences) 50µM was added to the intracellular solution above to eliminate voltage-dependent sodium currents. The same laser stimulation parameters were used to stimulate labeled ChR-2 expressing fibers and terminals. ChR-2 excitation is typically performed using blue light; however the dichroic mirror necessary for use of a 488 nm laser does not allow for visualization of the red fluorescent latex beads used to visualize back-labeled corticocollicular neurons. Therefore a UV laser was used, which we (and others (Petrof et al., 2015)) have found to robustly activate ChR-2, given the extended short-wavelength tail of the ChR-2 absorption spectrum (Zhang et al., 2007).

### Laser Map Analysis

Electrophysiological analyses were performed with Clampfit followed by further analysis in custom written MATLAB software. For each pixel, a pre-stimulation baseline period of 100 msec was compared to a post-stimulation analysis period of 150 msec. Statistical maps were created by computing the F-statistic (variance of the post-stimulation analysis period divided by the variance of the baseline period) for each stimulation site and maps were thresholded based on a ratio of 2.0. F-statistics were used to account for pre-stimulation variability and spontaneous activity, and are commonly used in the brain imaging literature (Howseman et al., 1997; Friston, 1998; Bowman, 2014). For quantitative comparison of inward or outward currents, currents were integrated over 150 msec to compute the total inward or outward charge transfer. To minimize the impact of spontaneous inward or outward currents, sites were only counted if two or more adjacent sites produced inward or outward currents in the recorded neuron, as previously described (Sturm et al., 2017). Identical 30 × 30 stimulation grids were used for each neuron. Therefore, to combine data across neurons, maps were aligned to the recording site. A 4 × 4 Gaussian kernel with a standard deviation of 0.5 pixels was convolved across each map to minimize the impact of subtle misalignment, as is ubiquitously done in the imaging literature (Worsley et al., 2002; Penny et al., 2011; Gramfort et al., 2015), and maps were averaged, pixel-by-pixel, across neurons. For display only, heatmaps were smoothed using bilinear interpolation in MATLAB.

### Statistical analysis

All statistical tests were run using MATLAB or SPSS. Shapiro-Wilk testing for normality was performed on all data sets, and in all cases data were found to be non-normal. Therefore, non-parametric testing was done. Specific tests (Mann-Whitney U-test, Spearman’s correlation, Chi Square and Kolmogorov-Smirnov 2-sample test) were applied as described in the Results section. To account for multiple comparisons, a post-hoc Holm-Bonferroni approach (Holm, 1979) was used.

## Results

### Mice and injections

Seven adult (30-60 days of age) BALB/c mice were injected with red retrobeads into the inferior colliculus to backlabel layer 5 and layer 6 corticocollicular cells (Figure 1A-C). An additional five mice had a combination of red retrobeads injected into the inferior colliculus and AAV-CaMKIIa-hChR2(H134R)-mCherry into the auditory thalamus (5 animals, Figure 1D-F) to permit thalamic stimulation of identified corticocollicular neurons.

### Cell-attached recordings

To determine the functional spatial resolution of the laser stimulation parameters described below in the Methods, excitation profiles in cortical neurons were measured using cell-attached recordings from unlabeled neurons in layers two through six (Layer 6: n = 3, Layer 5: n=4, Layer 4: n=5, Layer 2/3: n= 4), similar to previous studies (Zarrinpar and Callaway, 2006; Sturm et al., 2014; Kratz and Manis, 2015). A 10×10 spot grid, with 10 µm spacing between adjacent rows and columns, was centered over the recorded cell to direct UV laser photostimulation (Figure 1G). Four mapping trials were run to measure the spiking reliability of auditory cortical neurons to a given stimulation point. Stimulation locations in which the cell responded with a spike (inset Figure 1G) in half or more trials were considered reliably driven and therefore used for analysis. All cells sampled (n=16) had relatively small excitation profiles (example shown in Figure 1H and 1I). Reliable photostimulation-driven spiking (i.e., seen on at least 50% of the trials) was observed with a mean width of 20.0 ± 7.7 µm (S.D.) in the rostro-caudal direction and 30.9 ± 14.6 µm (S.D.) in the dorso-ventral direction (Figure 1J). This information was used to design later mapping experiments using a larger stimulus grid, for which 25 µm interstimulus spacing was used.

Corticocollicular neurons are found in layer 5 and layer 6 of the cortex, with the majority of the projection (~80% of backlabeled neurons) originating from layer 5. We found previously that the biophysical and morphological properties of these two cell types differ substantially (Slater et al., 2013), and as such, we hypothesized that their inputs would also differ. To test this hypothesis, sequential recording of paired pre-identified layer 5 and layer 6 corticocollicular neurons within 150 µm rostro-caudal distance of each other was done (n=8 cells in each layer). Laser photostimulation across 900 stimulus sites (30 × 30 grid) was used to uncage glutamate to measure local cortical excitatory inputs (by holding the cell at −60 mV) and inhibitory inputs (by holding the cell at +10 mV).

### Separation of direct and synaptic responses

Traditionally, a time window, typically in the range of 7-12 msec, is used to separate inward currents caused by direct activation of post-synaptic glutamate receptors on the recorded cell from synaptic responses induced by stimulation of cells providing synaptic input to the recorded cells (Jin et al., 2006; Hooks et al., 2013; Sturm et al., 2014; Meng et al., 2015). Using this time window-based approach, sites with latencies less than the predefined time window are deemed “direct” and sites with latencies beyond this window are deemed “synaptic.” One result of this approach is that stimulation sites near the recorded cell that produce a mixture of direct and synaptic inward currents are removed from the analysis, potentially biasing the inward current maps to only reflect more distant inputs. To remove this bias, similar to previous work (Staiger et al., 1999; Llano and Sherman, 2009), we generated repeat maps created in a synaptic blockade medium with low calcium (0.01 mM) and high magnesium (4.0 mM) and subtracted the synaptic blockade map (which should reflect only direct currents) from the total inward current map. The resulting subtraction map should only reflect synaptic currents. An example for an identified layer 6 corticocollicular cell is shown in Figures 2A-D, showing the total inward current map (Figure 2A), the map obtained with low calcium aCSF (Figure 2B) and the map generated by subtracting the low calcium map from the normal aCSF map (Figure 2C). As shown in Figure 2C, most of the synaptic input to this cell is from nearby regions in layer 6. An example of a site that would have been eliminated using the traditional time window method, but retained using the low-calcium method, is shown in Figure 2D (corresponding to the asterisk in Figures 2A-C). After glutamate stimulation, this site produced two inward currents: an early inward current that was retained after synaptic blockade, and a later current, starting approximately 9 msec after the direct current, which was presumably synaptic in nature. These data suggest that the low calcium method retains activation sites that would have been eliminated using the time window method.

**Figure 2:**
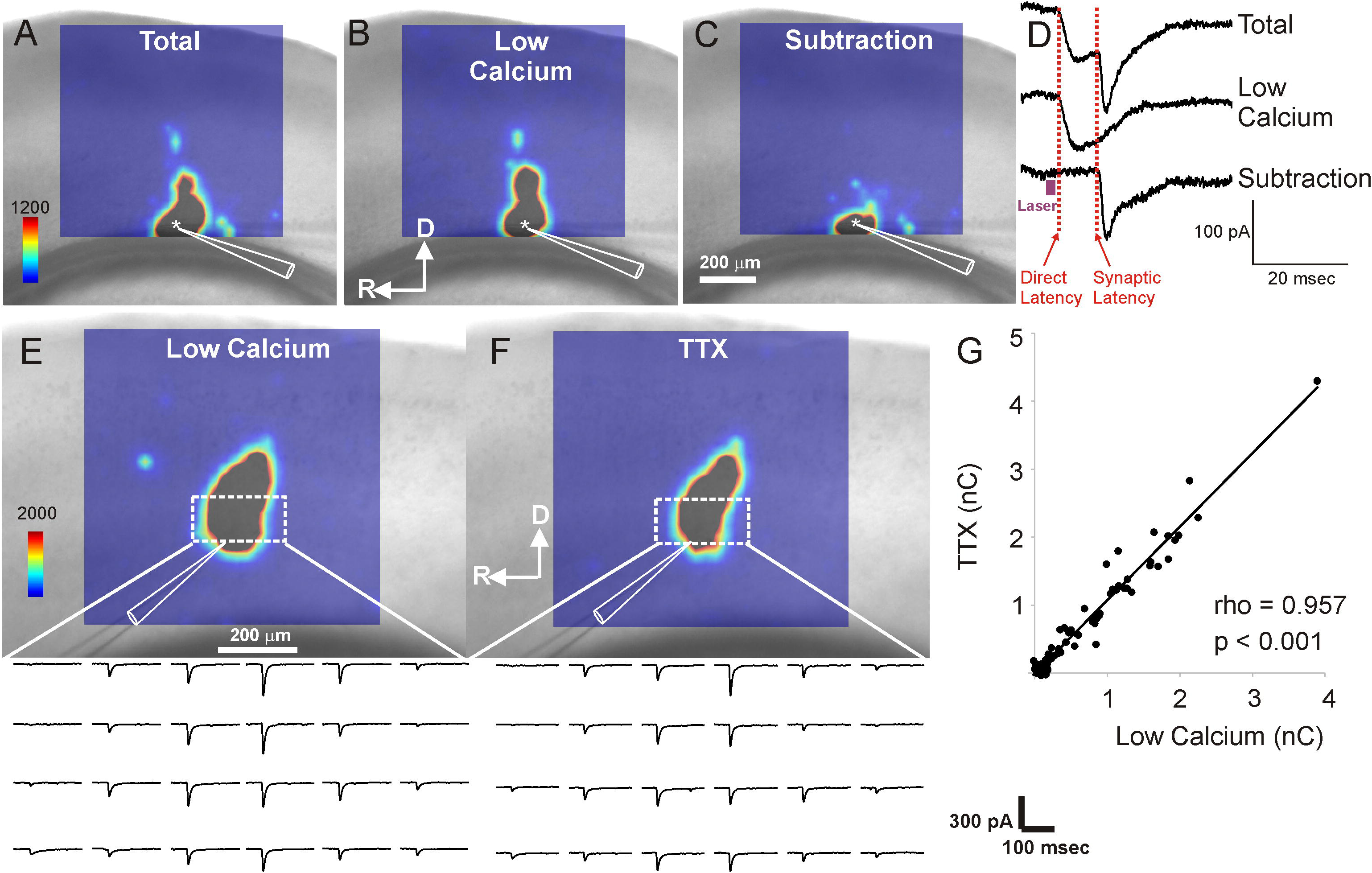
Low calcium technique to separate direct vs. synaptic inputs. (A) Total inward current map in a layer 6 corticocollicular neuron. (B) Same neuron, but here the map is shown using inward currents obtained using low calcium aCSF. (C) Subtraction map, obtained by subtracting the individual traces used to obtain the map shown in (B) from the current traces used to create the map shown in (A). (D) Traces taken from the laser stimulation site denoted with an asterisk, demonstrating that this site contains a short-latency, multipeaked response, with the latter portion of the response being eliminated using a low-calcium aCSF solution, suggesting that this latter portion is synaptic in nature. (E) Direct inward current map of a layer 5 neuron obtained using low-calcium aCSF. Below the map is a series of current traces taken from the boxed area of the map. (F) Direct inward current map of a layer 5 neuron obtained using TTX. Below the map is a series of current traces taken from the boxed area of the map. (G) Correlation of the amplitude of responses of the direct inward current obtained using low calcium aCSF (abscissa) and TTX (ordinate), measured at each stimulation site eliciting a response. Correlation coefficient (0.957) obtained using Spearman’s correlation. Triangular outlined image corresponds to a diagram of the recording pipette, the tip of which points to the location of the recorded cell.

Although the gold standard for eliminating synaptic inputs is the use of TTX (Callaway and Katz, 1993; Sturm et al., 2017), the long duration of action of TTX makes it difficult to record from more than one cell in a slice. To determine if the map obtained under TTX (dissolved in otherwise normal aCSF) is similar to the map obtained under low-calcium conditions, a layer 5 neuron was recorded under both conditions, and shown in Figures 2E-G. Figure 2E shows the inward current map of the cell under low calcium conditions while Figure 2F shows the same cell exposed to TTX. As shown, their maps are qualitatively similar. Correlation of the inward currents generated under the two conditions is shown in Figure 2G. As shown, the inward currents are highly correlated (Spearman’s rho = 0.954, p < 0.001). These data suggest that a low-calcium solution mimics data obtained under TTX. Therefore, for analysis of local input maps described below, three maps were obtained: a map obtained in routine aCSF, with cells held at −60 mV, a map obtained in low calcium aCSF, with cells held at −60 mV, and a map obtained in routine aCSF with cells held at +10 mV to record inhibitory inputs.

### Laser photostimulation mapping of inputs to identified layer 5 and layer 6 corticocollicular cells

Using the approach described above, we recorded from eight non-sequentially recorded pairs of neurons: one identified layer 5 corticocollicular cell and one identified layer 6 corticocollicular cell. An example of such a pair of recordings is shown in Figure 3. As shown in Figure 3A and B, the layer 5 corticocollicular neuron receives prominent synaptic input from a region near the cell body in layer 5 as well as from a region approximately vertically aligned with the cell body in layer 2/3. The excitation and inhibition appear to be roughly spatially matched. Synaptic traces from representative laser stimulation sites show mostly short-latency, multi-peaked induced currents.

**Figure 3:**
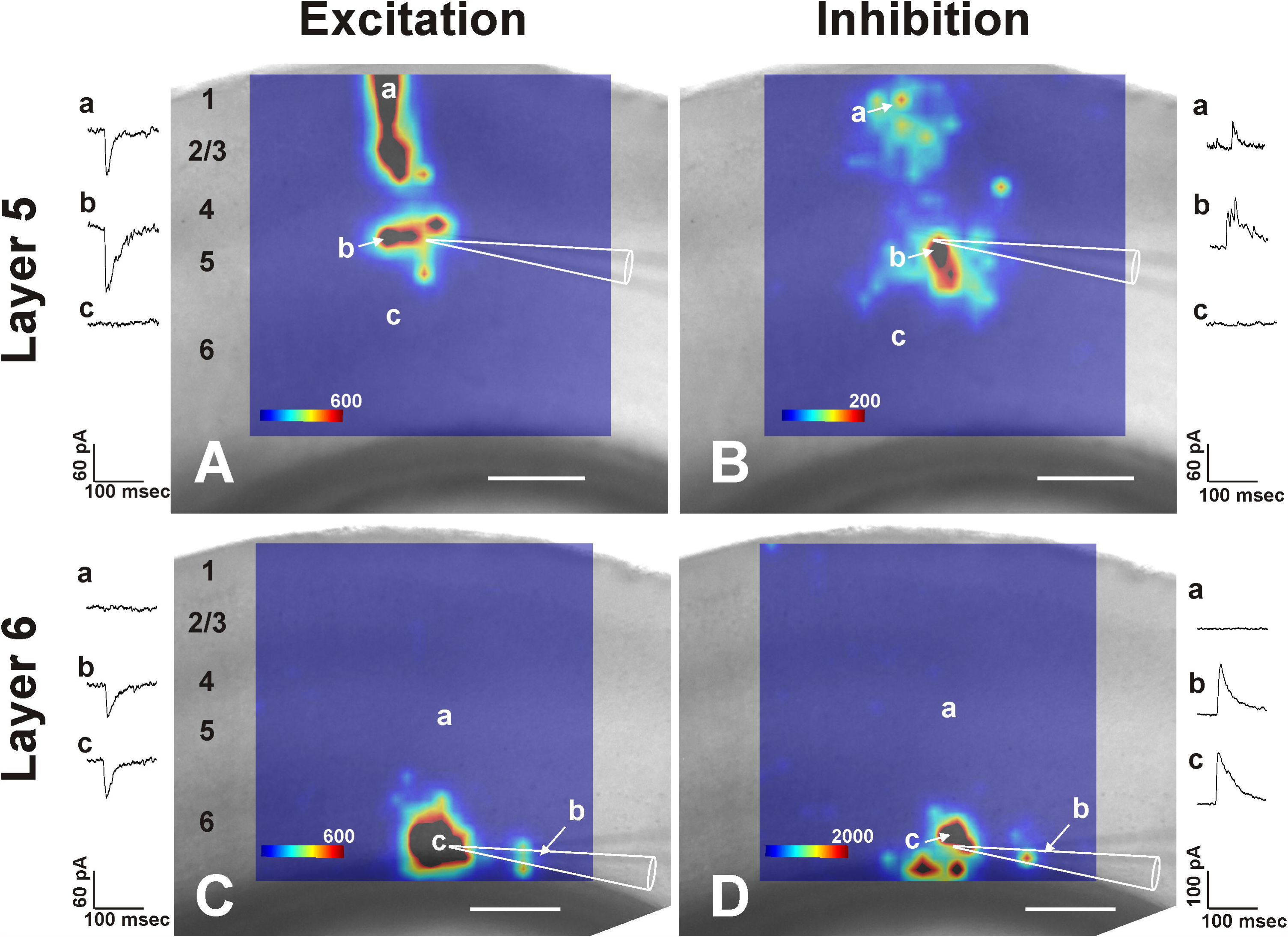
Example of synaptic input maps from matched layer 5 and layer 6 corticocollicular neurons. (A) Excitatory synaptic input map from a layer 5 corticocollicular neuron. (B) Inhibitory synaptic input map from a layer 5 corticocollicular neuron. (C) Excitatory and (D) inhibitory synaptic input maps from a layer 6 corticocollicular neuron taken from just below the layer 5 neuron in (A) and (B). In all cases, traces from individual stimulation sites shown in a, b and c, which correspond to locations denoted on the adjacent heat maps. Scale bar in all images = 200 μm.

In contrast, a layer 6 corticocollicular cell immediately ventral to the layer 5 corticocollicular cell described above has a much more spatially restricted synaptic input area. As shown in Figures 3C and D, this cell primarily receives input from layer 6. This restricted degree of input to this layer 6 cell is unlikely to be related to poor slice connectivity or poor slice health since the layer 5 cell from the same slice retained excellent connectivity, and the synaptic currents recorded in the layer 6 corticocollicular cell were as robust as those recorded from the layer 5 corticocollicular cell from the same slice (see synaptic traces adjacent to each map).

### Composite laser stimulation maps

Maps across all layer 5 corticocollicular cells and layer 6 corticocollicular cells were aligned and averaged to produce composite maps for layer 5 corticocollicular cell synaptic inward currents (Figure 4A, top left), layer 5 corticocollicular cell outward currents (Figure 4A, top right), layer 6 corticocollicular cell synaptic inward currents (Figure 4A, bottom left) and layer 6 corticocollicular cell outward currents (Figure 4A, bottom right). Currents were collapsed across the vertical or the horizontal dimension to produce plots of synaptic input across the rostro-caudal dimension (shown above or below the maps) or across the dorso-ventral dimension (shown to the left or right of the maps).

**Figure 4:**
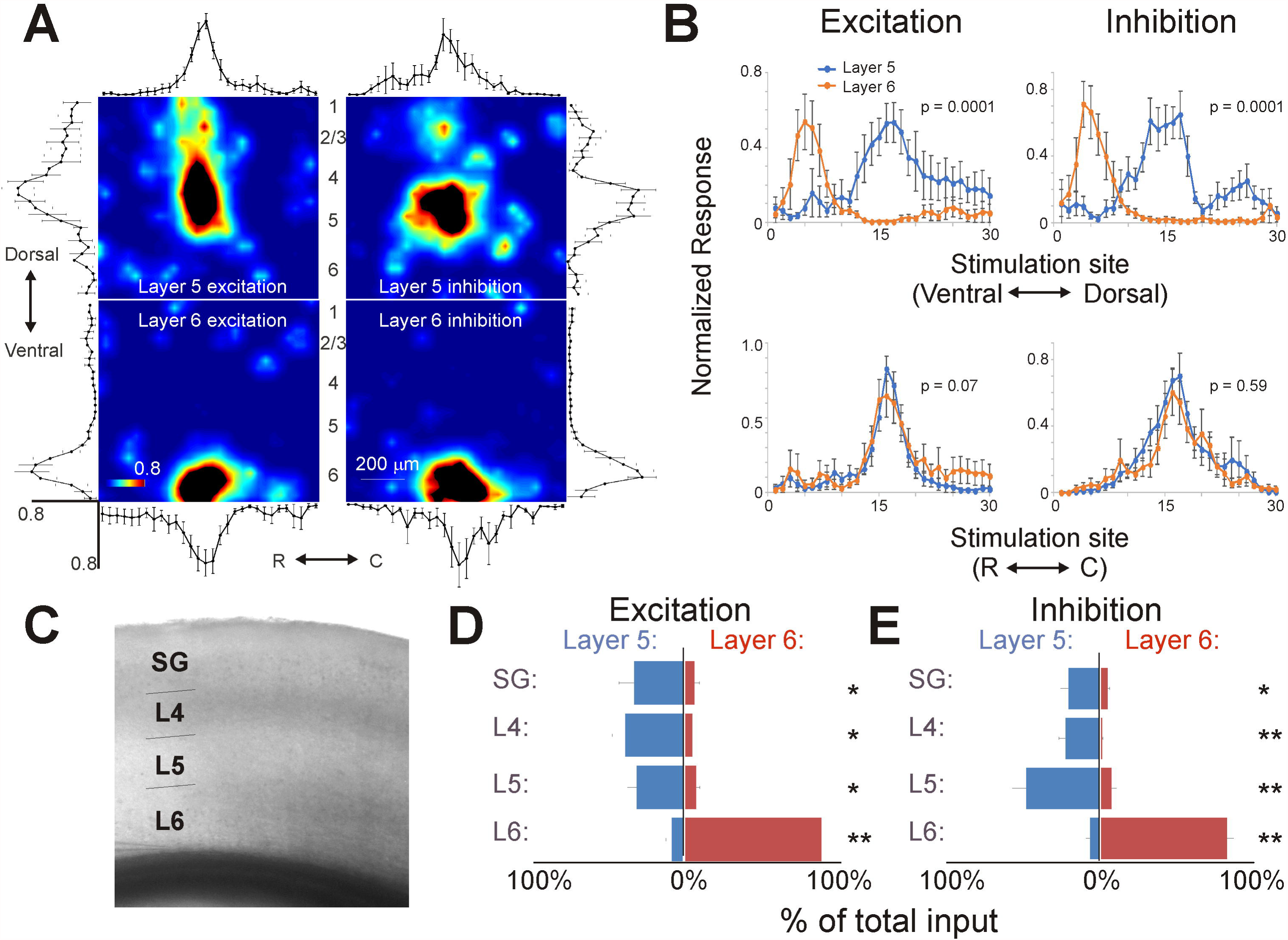
Summary of results comparing local excitation to layer 5 and layer 6 corticocollicular neurons. (A) Average of coregistered heatmaps for layer 5 (top) and layer 6 (bottom) corticocollicular neurons for both excitation (left) and inhibition (right). Shown above or below each heatmap are the average of the synaptic input, collapsed across all rows. Shown to the left or right of each heatmap are the average of the synaptic input magnitudes, collapsed across all columns. In each case, the average synaptic input for each column or row is expressed as a normalized value relative to the maximum synaptic input to that neuron. Error bars = standard errors. (B) The same average synaptic input maps collapsed across columns (top) and rows (bottom) shown in (A), replotted to permit direct comparison of layer 5 vs. layer 6 corticocollicular neurons. Distance along abscissa corresponds to the 30 rows (top) or columns (bottom) of stimulation sites. p values were generated using the Kolmogorov-Smirnov 2-sample test. (C) Representative raw image showing the delineation of the layers (SG=supragranular, L4=layer 4, L5=layer 5, L6=layer 6) used for the analyses in (D) and (E). (D) and (E) Two butterfly graphs comparing the mean excitatory (D) and inhibitory (E) input from individual layers to layer 5 and layer 6 corticocollicular neurons. *p<0.05, **p<0.01, comparisons made with Mann-Whitney U-test.

Qualitatively, the inward and outward current maps of layer 5 corticocollicular neurons differ substantially from those derived from layer 6 corticocollicular neurons because of the prominent excitatory and inhibitory input from layer 2/3 onto layer 5 corticocollicular neurons. An analogous set of apical inputs is not seen in layer 6 corticocollicular neurons. This difference is reflected in the differences in the plots of synaptic input seen across the dorso-ventral dimension (Figure 4B, top row), which were compared using a 2-sample Kolmogorov-Smirnov test, which revealed a highly significant difference in terms of the distribution of currents generated across stimulation sites (p = 0.0001 for excitation and p = 0.0001 for inhibition). In contrast, no statistically significant differences were seen across the rostro-caudal axis (p = 0.07 for excitation p = 0.59 for inhibition, Figure 4B, bottom row). These data suggest that layer 5 and layer 6 corticocollicular neurons integrate synaptic inputs from different layers of cortex, but that they integrate from similar regions across the rostro-caudal axis.

To confirm that the differences seen in the dorso-ventral dimension conform to different layers of cortex, synaptic inputs were pooled within layers, based on high-contrast images of the cortex to identify each layer (for example image, see Figure 4C). Inputs from layers 1-3 were pooled and referred to as “supragranular (SG)”. Direct comparison between inputs from any given layer also revealed significant differences between layers (Figure 4D). For example, compared to layer 5 corticocollicular neurons, layer 6 corticocollicular neurons showed significantly more excitatory and inhibitory input from layer 6, (p = 0.002 and p = 0.0009, respectively). In contrast, layer 5 corticocollicular neurons received greater excitatory and inhibitory input from all other cortical layers (Layer 5 excitatory input: p = 0.01, Layer 5 inhibitory input: p = 0.002, Layer 4 excitatory input: p = 0.016, Layer 4 inhibitory input: p = 0.003, SG excitatory input: p = 0.041, SG inhibitory input: p = 0.024). These data suggest that layer 5 corticocollicular neurons vertically integrate excitatory and inhibitory inputs from the pia to layer 5, while layer 6 corticocollicular neurons integrate inputs primarily from layer 6.

### Assessment of direct thalamic inputs to corticocollicular neurons

Recent work has shown that neurons in both layer 5 and layer 6 receive direct input from the thalamus (Viaene et al., 2011; Constantinople and Bruno, 2013; Yang et al., 2014; Ji et al., 2015). The presence of a direct thalamic input to corticocollicular neurons would significantly impact models of corticofugal function since such an input would provide a short-latency mechanism to rapidly modulate ascending information. To determine if layer 5 or layer 6 corticocollicular neurons received direct thalamic input, we initially used glutamate uncaging in the auditory thalamus, but found relatively sparse connectivity in the adult animal in 300 μm slices. In addition, this approach cannot distinguish between mono- and polysynaptic thalamic input to the recorded cortical neurons. Therefore, virally-mediated expression of channelrhodopsin-2 (ChR-2) was used to label thalamocortical fibers. AAV-CaMKIIa-hChR2(H134R)-mCherry virus (AAV2) was injected into the MGB to label the thalamic afferents (See Figure 5A for a diagram of the double-injection animal preparation). ChR-2 has been utilized previously in this manner to map long range projections (Petreanu et al., 2007; Petreanu et al., 2009; Cruikshank et al., 2010).

**Figure 5:**
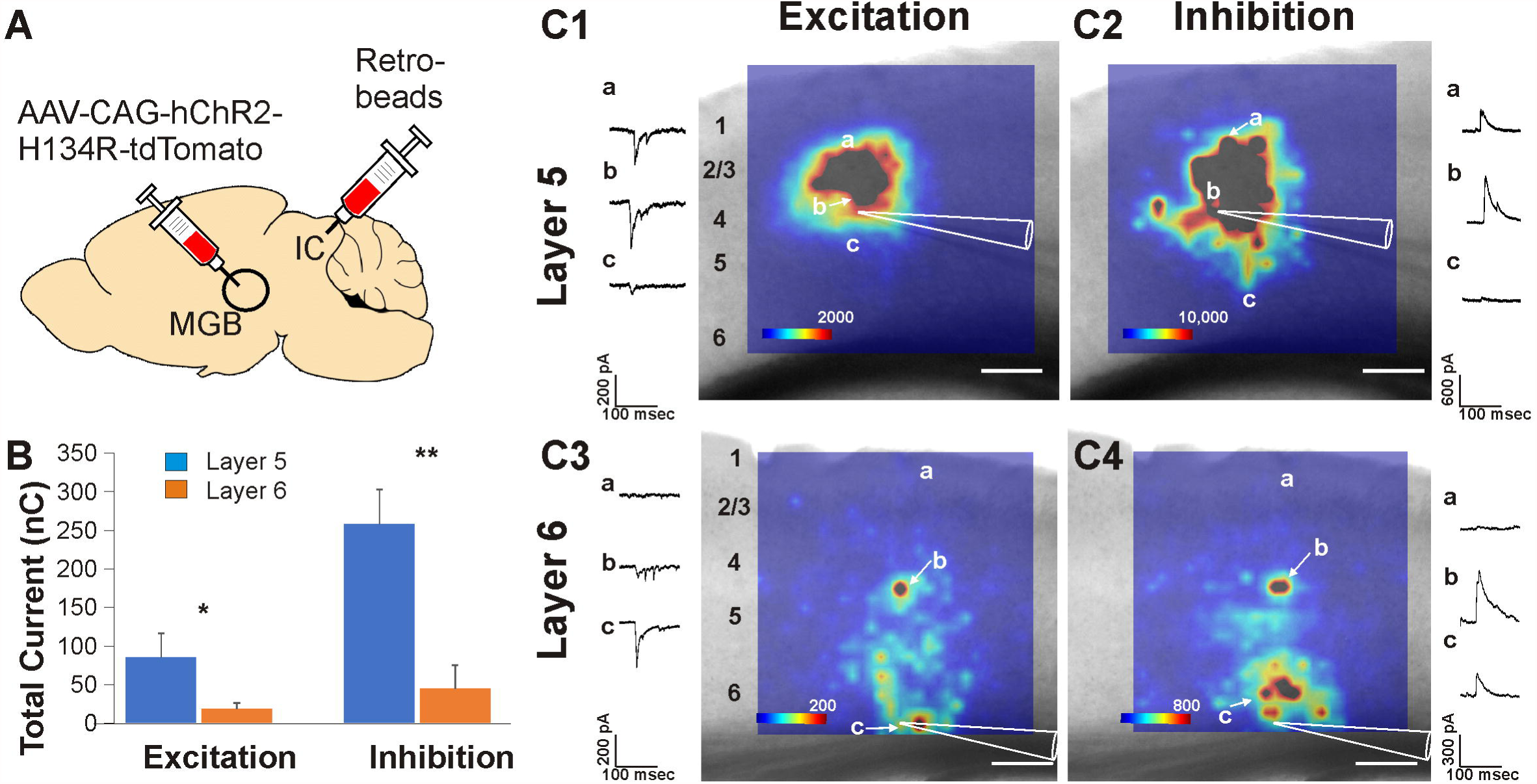
Mapping thalamic inputs to corticocollicular neurons. (A) Schematic diagram illustrating the dual injections of an AAV virus expressing ChR-2 in the MGB and red retro-beads into the inferior colliculus (IC). (B) Average excitatory and inhibitory current across all recorded cells (n=5 per layer) elicited by stimulation of thalamocortical afferents. *p<0.05, **p<0.01, comparisons made with Mann-Whitney U-test. (C1) Excitatory thalamic input map from a layer 5 corticocollicular neuron. (C2) Inhibitory di-(or poly-) synaptic thalamic input map from a layer 5 corticocollicular neuron. (C3) Excitatory thalamic input map from a layer 6 corticocollicular neuron. (C4) Inhibitory di-(or poly-) synaptic thalamic input map from a layer 6 corticocollicular neuron. In all cases, traces from individual stimulation sites shown in a, b and c, which correspond to locations denoted on the adjacent heat maps. Scale bar in all images = 200 μm.

Thalamic input responses were recorded from retrogradely labeled corticocollicular neurons and were mapped in both layer 5 and 6 corticocollicular neurons (n=5 in each layer) using a 30×30 grid that covered much of the cortex as with the glutamate uncaging experiments. Both excitatory and inhibitory currents were measured. As previously noted in other studies of thalamocortical afferents (Cruikshank et al., 2010; Liu et al., 2011; Ji et al., 2015), inhibitory inputs were prominent. Inhibition was larger in layer 5 corticocollicular neurons compared to layer 6 corticocollicular neurons (260 ± 90 nC versus 45.7 ± 60 nC, p=0.003, Mann-Whitney U test). Excitation was also larger in layer 5 corticocollicular neurons 86.4 ± 62 nC compared to 19.2 ± 15 nC in layer 6 corticocollicular neurons (p=0.02, Mann-Whitney U test, see Figure 5B). Layer 5 corticocollicular neurons received excitatory and di-(or poly-) synaptic thalamic input from a broad region from layer 5 and extending to layer 2/3 (see Figures 5C1 and C2 for examples). Layer 6 corticocollicular neurons received synaptic input from thalamocortical axons in layer 6. In addition, unlike the local synaptic input maps seen with caged glutamate, layer 6 corticocollicular neurons received an additional synaptic input from thalamocortical axons located at the border of layers 4 and 5 (for examples, see Figures 5C3 and C4).

From these data, it is apparent that both layers of interest receive either direct or indirect thalamic input. However, it is unclear from these experiments whether these thalamic inputs are mono- or poly-synaptic. Initially, to assess whether layer 5 and layer 6 corticocollicular neurons receive a monosynaptic thalamic input, the data generated from the mapping experiments were used to measure latency and jitter as indicators of a monosynaptic response (Doyle and Andresen, 2001; Karayannis et al., 2006). From the 30×30 grid of points used to stimulate thalamocortical afferents, a 3×3 sub-grid of adjacent points around the cell was used to generate latency and jitter information from each of the cells sampled. Latency was measured as the time between stimulation and ten percent of the peak rise time of the current. Jitter was calculated as the standard deviation of the latency.

It was found that both layer 5 (n = 5) and layer 6 (n = 5) corticocollicular neurons receive input from ChR-2 labeled fibers shortly after stimulation. Layer 5 corticocollicular neurons had shorter latencies with an average of 2.66 ± 0.35 msec versus layer 6 corticocollicular neurons which had an average of 3.04 ± 0.22 msec (n=5 in each layer, p = 0.048, Mann-Whitney U test). Similarly, the jitter of the response was smaller in layer 5 corticocollicular neurons (0.31 ± 0.22 msec) compared to layer 6 corticocollicular neurons (0.90 ± 0.42 msec, n = 5 in each layer, p = 0.017, Mann-Whitney U test, see Figures 6A and 8B).

**Figure 6:**
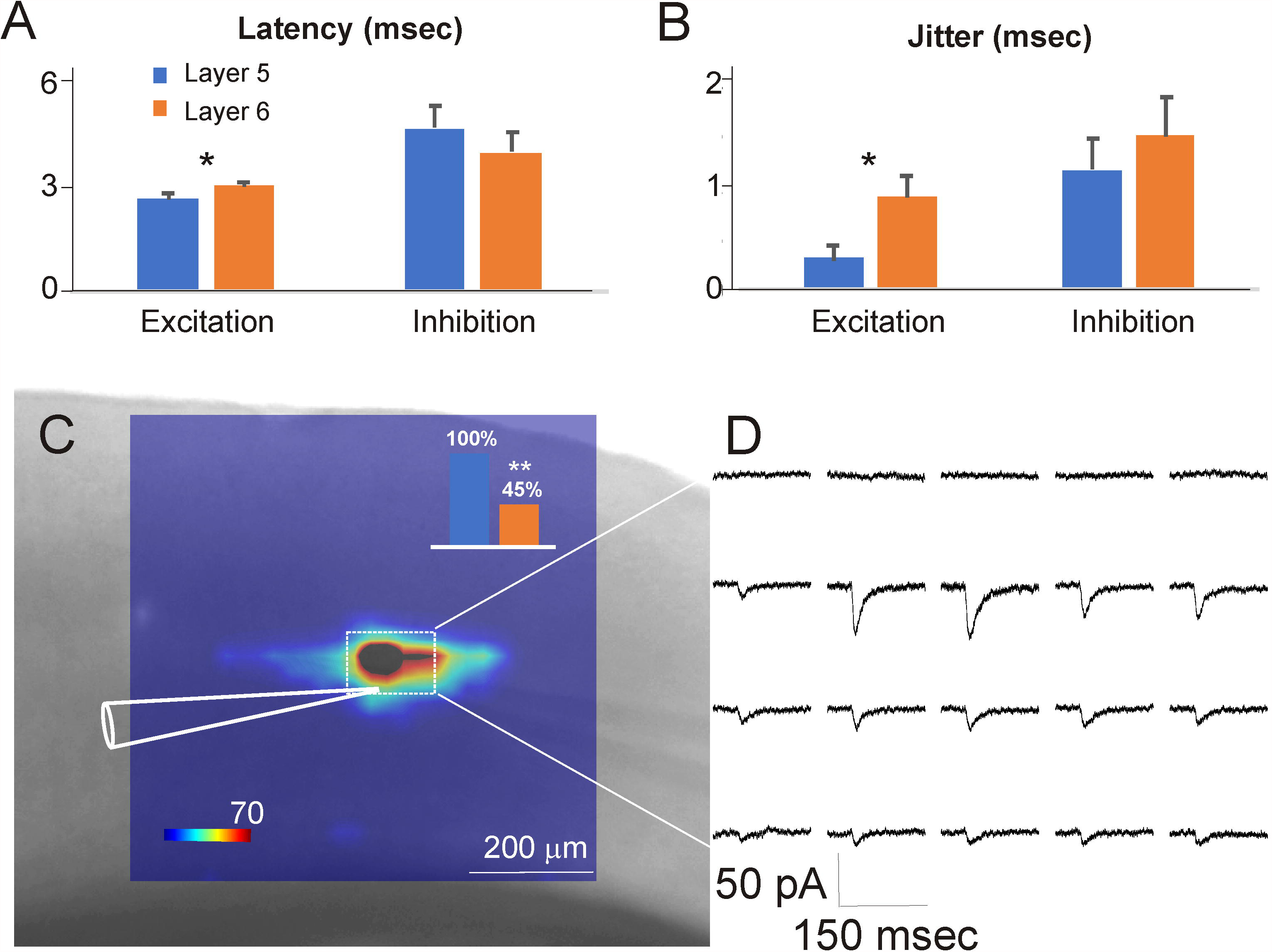
Direct inputs from the thalamus to corticocollicular neurons. (A) Average latency from excitatory and inhibitory after stimulation of labeled thalamocortical axons to layer 5 and layer 6 corticocollicular neurons. (B) Average jitter from excitatory and inhibitory after stimulation of labeled thalamocortical axons to layer 5 and layer 6 corticocollicular neurons. *p<0.05 using Mann-Whitney U test. (C) An example of an excitatory heat map in a layer 5 corticocollicular neuron obtained in 2 µM TTX. Inset shows percentage of layer 5 corticocollicular neurons (13/13 or 100%) and layer 6 corticocollicular neurons (5/11) showing monosynaptic responses. **p<0.005 using Chi-Square test. (D) Example traces from the boxed area in (C).

Latencies and jitters were also measured for inhibitory responses, which are assumed to be at least disynaptic since thalamocortical terminals release glutamate and are therefore excitatory (Kharazia and Weinberg, 1994). Inhibitory latencies were significantly longer in layer 5 corticocollicular neurons at 4.77 ± 1.20 msec compared to excitatory latencies which were 2.66 ± 0.35 msec (n = 5, p = 0.0079, Mann-Whitney U test). Layer 5 also had larger jitter (1.16 ± 0.60 msec) for inhibitory responses versus 0.31 ± 0.22 msec for excitatory responses (n=5, p = 0.016, Mann-Whitney U test). Layer 6 corticocollicular neurons, similarly, trended to have longer latencies for inhibitory responses 4.01 ± 1.13 msec, compared to excitatory responses 3.04 ± 0.22 msec (n=5, p = 0.095, Mann-Whitney U test). There was a nonsignificant trend for the jitter in layer 6 corticocollicular neurons to be larger for inhibitory currents: 1.48 ± 0.76 msec for inhibitory compared to 0.90 ± 042 msec for excitatory currents (n=5, p = 0.18 Mann-Whitney U test, see Figures 6A and 6B).

To unambiguously address whether layer 5 and layer 6 corticocollicular neurons receive monosynaptic inputs from the thalamus, in a subset of experiments 2µM TTX was added to the circulating aCSF to block voltage gated sodium channels. Activation of ChR-2 at the synapse will cause a release of neurotransmitter without requirement of the activation of voltage-gated sodium channels. Using TTX has been shown to limit polysynaptic propagation (Petreanu et al., 2009; Ji et al., 2015) by restricting input to only fibers in contact with a postsynaptic cell. The axonal potassium channel blocker 4-aminopyridine, which is often used to increase terminal release excitability in experiments involving TTX (Shu et al., 2007; Petreanu et al., 2009; Tritsch et al., 2012), was not necessary here given the strong laser-induced post-synaptic currents that were seen, likely due to the high release probability of the thalamocortical synapse (Gil et al., 1999).

Using this approach, it was found that optical stimulation of thalamocortical terminals produced excitatory post-synaptic currents at least 3SD above baseline noise calculated using the first 200 msec of the trace, suggesting direct synaptic contacts between these terminals and the recorded neurons. Layer 5 corticocollicular neurons receive direct input from the thalamus more often (11 out of 11 cells) than layer 6 corticocollicular neurons (5 out of 13 cells, χ^2^ = 8.25, p=0.004, see Figure 6C and D for an example of a layer 5 corticocollicular neuron receiving direct thalamic input). The introduction of TTX narrows the extent of input elicited with ChR-2 stimulation. Therefore, it is likely that the total excitatory thalamic input (i.e., before adding TTX) to corticocollicular neurons is reflective of both the local cortical input driven by the thalamus as an ensemble and the direct thalamocortical input to a given corticocollicular neuron. This type of organization would indicate that the corticocollicular pathway is a nexus for direct thalamic and local cortical inputs.

## Discussion

### Summary

In the current study, layer 5 and layer 6 corticocollicular neurons were found to receive distinct patterns of local cortical input such that layer 5 corticocollicular neurons receive significantly more excitatory and inhibitory synaptic input from layer 5 and upper cortical layers, while layer 6 corticocollicular neurons receive a greater degree of corresponding input from layer 6. As part of this study, a low calcium trace subtraction approach to separate synaptic from direct glutamate-induced inward currents was validated. In addition, it was determined that all recorded layer 5 corticocollicular cells receive direct input from the auditory thalamus, while a minority of layer 6 corticocollicular cells receives direct thalamic input. The implications of these findings are discussed below.

### Technical considerations

The time window method is a commonly used method to separate direct versus synaptic stimulation induced by the uncaging of glutamate (Jin et al., 2006; Hooks et al., 2013; Sturm et al., 2014; Meng et al., 2015). A major practical point in favor of this technique is that it requires no extra caged glutamate, relying on only one uncaging map to yield results. However, others have shown that direct inputs to distal dendrites can fall outside of the window commonly used to separate direct and synaptic input, and conversely that synaptic inputs close or directly on the soma can fall within the window (Petreanu et al., 2009). In the current study, sites near the soma showed a mixture of direct and synaptic responses (Figure 2D) that would normally be eliminated using the time-window technique. Validation using TTX shows close correspondence between the TTX and the low calcium trace subtraction maps (Figures 2E-G). The bias of the time window method to emphasize distal inputs may explain why excitatory input maps in this and previous work (Llano and Sherman, 2009), generally show less relative distal (layer 2/3) input onto recorded layer 5 neurons than previous studies (Schubert et al., 2001; Schubert et al., 2006; Jacob et al., 2012).

### Circuit and functional implications

Layer 5 and layer 6 corticocollicular neurons were found to be embedded in different local cortical circuits, which may influence how they modulate the inferior colliculus. Layer 5 corticocollicular cells share many properties with layer 5 corticothalamic cells: i) both populations comprise large pyramidal cells with tufted thick apical dendrites, ii) both demonstrate intrinsic bursting firing patterns and iii) the current study suggests that both receive substantial excitatory and inhibitory input from layers 2-5 (Llano and Sherman, 2009). Both sets of cells also express the retinol binding protein-4 marker (Gerfen et al., 2013; Xiong et al., 2015). Given prior work in other sensory and motor systems showing branching from layer 5 neurons to thalamus and other subcortical structures (Deschênes et al., 1994; Bourassa et al., 1995; Kita and Kita, 2012) these data raise the possibility that at least a subset of layer 5 auditory corticocollicular neurons send a branch to the thalamus. Layer 5 auditory cortical neurons also project to other subcortical targets such as the corpus striatum (where there is evidence for branching from corticocollicular neurons (Moriizumi and Hattori, 1991)), and cochlear nucleus (Doucet et al., 2003), raising the possibility that branches to multiple subcortical structures exist in the layer 5 auditory corticofugal system. In contrast, the layer 6 corticocollicular neuronal morphological properties are unique within the cortex, having elongated and thin dendritic arborizations, in some cases reaching the pia and other cases extending into adjacent cortical columns (Slater et al., 2013). However, their excitatory synaptic inputs do not seem to differentiate them from other layer 6 neurons such as the corticothalamic neurons (Llano and Sherman, 2009). This finding is somewhat surprising given their extensive dendritic arborizations, which do not appear to receive input from local cortical neurons, and particularly from the upper layers. It is possible that coordinated input is required to stimulate these cells, as groups of cells are often activated in ensembles, particularly from thalamic inputs (Bruno and Sakmann, 2006; Wang et al., 2010). This supposition is supported by the current finding that optogenetic stimulation, which activates populations of afferent fibers, is better able to drive layer 6 corticocollicular neurons from distal sites than glutamate stimulation (e.g. compare Figures 5C3 and C4 to Figures 3C and D). As such, it is possible that focal glutamate stimulation is unable to activate the thin dendrites found in the layer 6 corticocollicular neurons in such a way that the activation is reflected in a somatic current.

### Thalamic connections to corticocollicular neurons

The findings in the current study that demonstrate direct connectivity between the thalamus and corticocollicular neurons are consistent with a growing body of literature that demonstrates thalamic inputs to layers other than layer 4 (Huang and Winer, 2000; Cruikshank et al., 2010; Viaene et al., 2011; Smith et al., 2012; Constantinople and Bruno, 2013; Krause et al., 2014; Ji et al., 2015). For example, a previous study in the somatosensory cortex found that pre-identified layer 6 corticothalamic neurons receive direct thalamic input (Yang et al., 2014). The investigators used electrical stimulation of thalamocortical afferents and found similar average latencies (2.52 msec) and jitter (0.17 msec) to our findings in layer 5, where the average latency = 2.41 msec and average jitter = 0.19 msec. The work presented here is the first evidence in the auditory system that there is a direct connection between the ascending projections and the descending projections, and the use of ChR-2 allowed for the definitive determination of monosynaptic thalamocortical input to corticocollicular neurons.

Combining the current data with previous work demonstrating differences in firing properties of layer 5 and layer 6 corticocollicular neurons (Slater et al., 2013), and work demonstrating that auditory corticocollicular neurons target the matrix portion of the lateral cortex of the IC while somatosensory inputs target GABAergic modules (Lesicko, Hristova et al. 2016), we propose that layer 5 cells receive short-latency thalamocortical input along with upper layer local cortical input, to rapidly send bursts of action potentials to neurons in the inferior colliculus. Layer 6 corticocollicular neurons are less likely to get direct thalamic input, but may respond to population thalamic activity, modulated by local inputs, to send trains of individual action potentials to neurons in the inferior colliculus (summarized in Figure 7). Given the numerous potential functions that have been hypothesized for the corticocollicular pathway (Zhang et al., 1997; Yan et al., 2005; Suga, 2008; Bajo et al., 2010; Xiong et al., 2015; Robinson et al., 2016), such a duplex system may be necessary to instantiate modulation that may require different time scales, different degrees of convergence or divergence, and different degrees of inhibition and excitation. Future work will elucidate the specifics of how these two pathways modulate the inferior colliculus.

**Figure 7:**
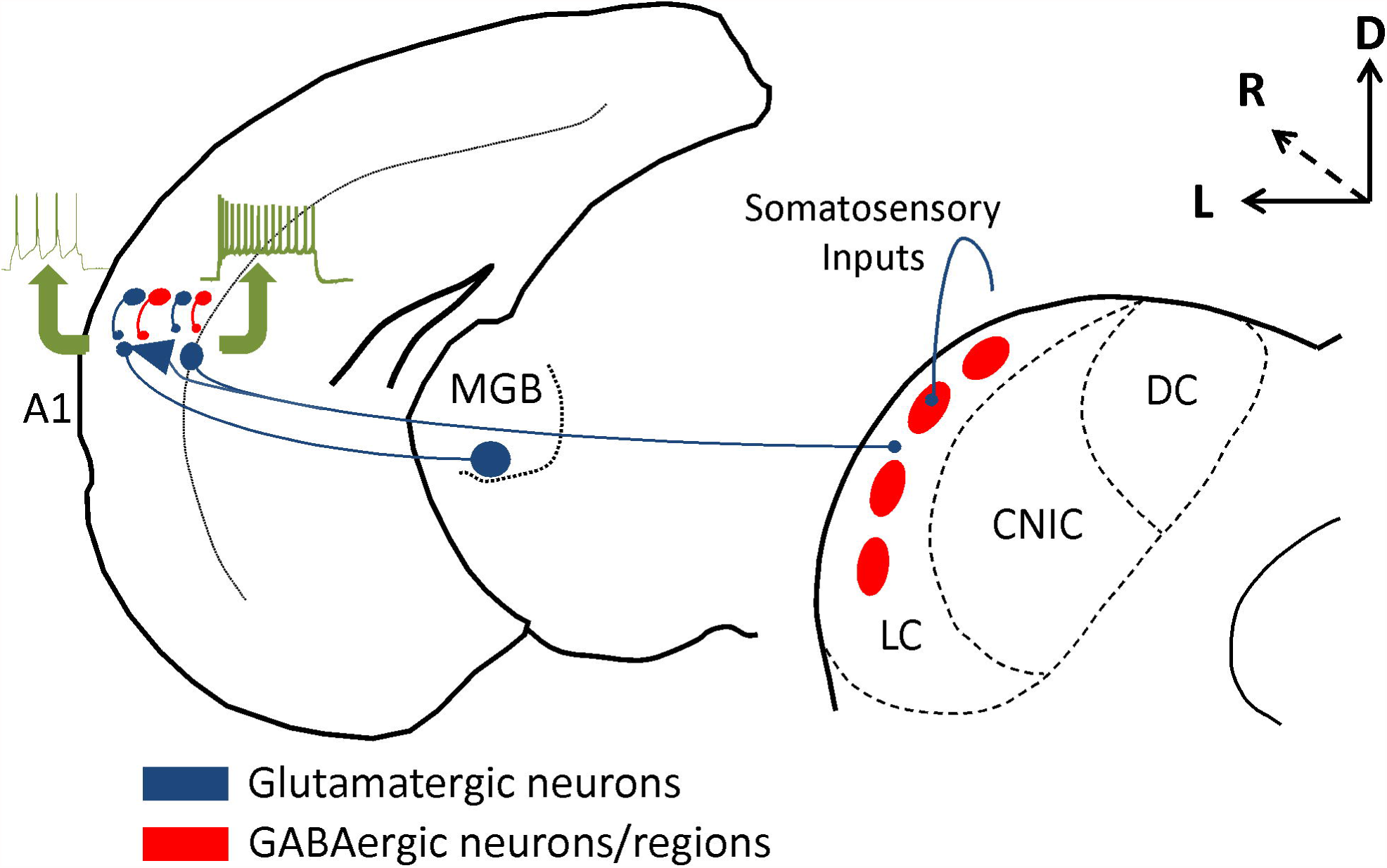
Updated model of auditory corticocollicular organization. Layer 5 corticocollicular neurons have intrinsic bursting properties (green arrow) and draw excitatory and inhibitory inputs primarily from upper and middle layers, while layer 6 corticocollicular neurons have regular spiking properties (green arrow) and draw excitatory and inhibitory inputs from layer 6. Firing properties are from (Slater et al., 2013). Layer 5 corticocollicular neurons receive a direct input from the MGB. Both sets of neurons project to the matrix portion of the LC while the GABAergic modules receive somatosensory inputs (Lesicko et al., 2016). A1 = primary auditory cortex.

## Acknowledgments

The authors thank Nhan Huynh for his MATLAB coding assistance. The authors also thank the University of Illinois Statistical Services for their assistance with the analysis.

